# A Biologically Plausible Mechanism to Learn Clusters of Neural Activity

**DOI:** 10.1101/389155

**Authors:** Adrianna R. Loback, Michael J. Berry

## Abstract

When correlations within a neural population are strong enough, neural activity in early visual areas is organized into a discrete set of clusters. Here, we show that a simple, biologically plausible circuit can learn and then readout in real-time the identity of experimentally measured clusters of retinal ganglion cell population activity. After learning, individual readout neurons develop *cluster tuning*, meaning that they respond strongly to any neural activity pattern in one cluster and weakly to all other inputs. Different readout neurons specialize for different clusters, and all input clusters can be learned, as long as the number of readout units is mildly larger than the number of input clusters. We argue that this operation can be repeated as signals flow up the cortical hierarchy.

---

Most areas of the brain represent information using populations of neurons, yet the coding scheme employed is still the subject of active investigation [1, 2]. One salient property of neural populations is the high redundancy with which they encode information. Redundancy is already high in the retina [3], and because of the great expansion in the number of neurons encoding sensory events, redundancy in the cortex is expected to be much higher [4–7]. In communications engineering, redundancy is viewed as valuable, because it allows for coding schemes that can correct errors introduced by transmission noise [8]. Redundant population codes also allow for downstream neural circuits to learn their statistics [4]. In fact, depending on the characteristics of neural noise, redundant population codes can be optimal for encoding information [9, 10]. In this context, an appealing principle for population codes is that correlations among neurons organize neural activity patterns into a discrete set of clusters, which can each be viewed as an error-robust population *codeword* [11–15].

The retinal ganglion cell (RGC) population has sufficient redundancy and/or correlation to organize neural activity into such clusters [13, 15]. Specifically, the probability landscape of retinal populations resembles a “mountain”, where the summit is the all-silent state, due to the sparseness of neural activity (Fig. 1) [11]. Radiating out from the summit in different directions in the space of neural activity patterns are a set of “ridges”. These ridges constitute *neural activity clusters* that are well-separated from each other and that can be identified as the latent states of a hidden Markov model [12]. Neural activity clusters have two other important properties. First, they exhibit error correction, meaning that when the same stimulus is repeated, one or few clusters are activated, despite the fact that the detailed neural activity pattern was often different on every trial. Second, clusters have receptive fields that are qualitatively different from those of their constituent ganglion cells, including some clusters with orientation selectivity [12].

**FIG. 1:**
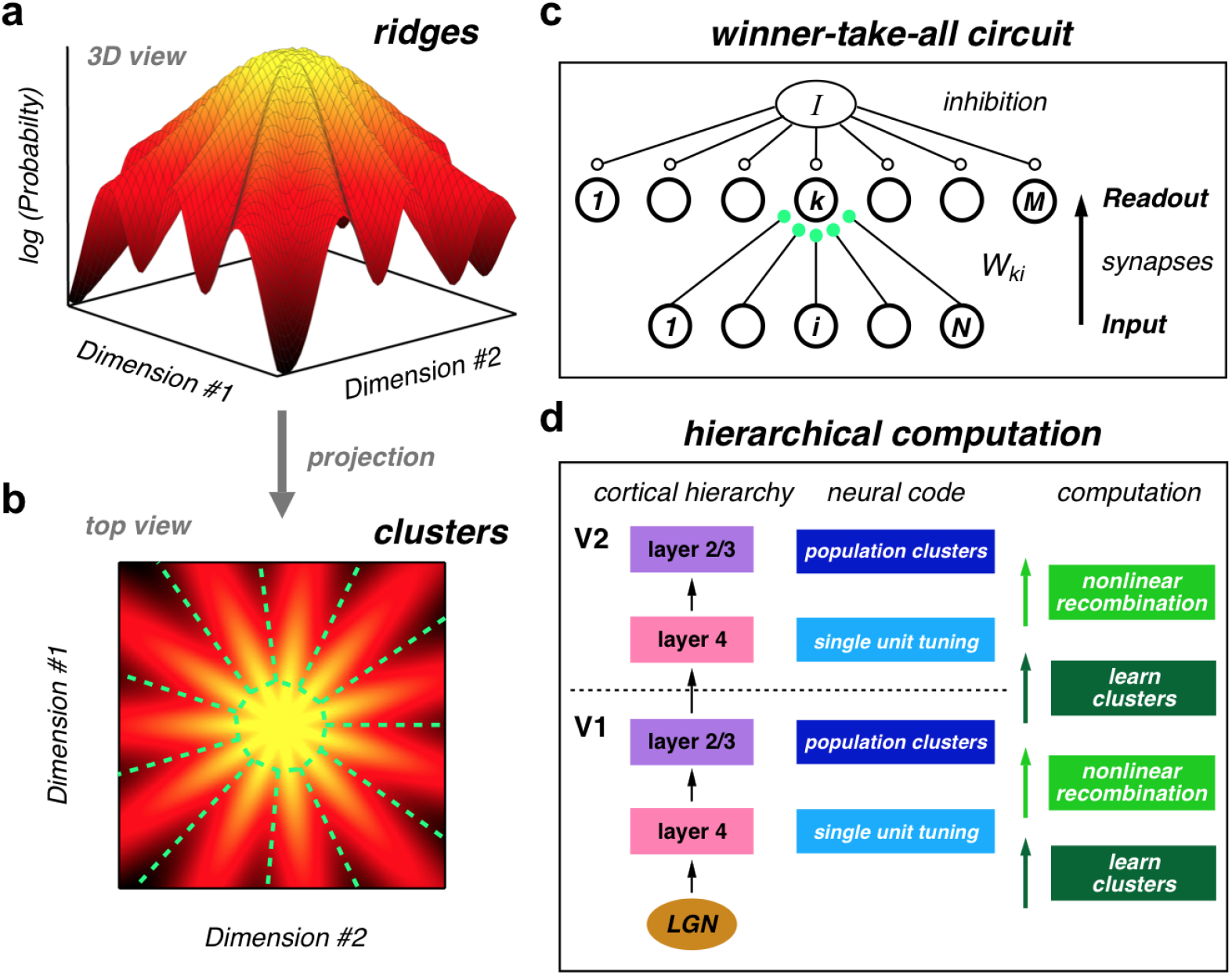
Learning clusters of population neural activity in a cortical hierarchy. (**a**) Schematic illustration of the global organization of population neural activity; the probability landscape is structured into a set of *ridges*. **(b)** Top view of the same probability landscape projected into population activity space; each ridge can be viewed as a cluster of neural activity (demarcated by dashed green lines). **(c)** Winner-take-all circuit to learn clusters. The input population of neurons *i* = 1, ⋯, *N* feeds into a layer of readout neurons, *k* = 1, ⋯, *M* with all-to-all synapses, *W_ki_*. Global inhibition, *I*, regulates the number of readout units to respond to each input. Hebbian and homeostatic plasticity of the feedforward synapses, *W_ki_*, learns clusters in real time. **(d)** Cluster learning is hypothesized to occur in layer 4 of the neocortical microcircuit. Layer 2/3 generates nonlinear recombinations of layer 4 neural activity, thus creating a new population code with new clusters of neural activity. These signals propagate up the cortical hierarchy, and the next layer 4 can again learn clusters in its input population, which are different than the lower-level clusters. This alternating process of cluster learning followed by recombination to form new clusters repeats going up the cortical hierarchy.

In previous work, we found that the strength of pairwise correlations between retinal ganglion cells was not precisely tuned to produce clusters, but was instead well within the range required [15]. The robustness of the clustered state suggests that neural populations elsewhere in the brain may also have sufficient correlation to be organized into clusters. This result makes it interesting to ask whether real neural circuits in the brain can learn such clusters? A compelling aspect of this idea is that downstream neurons that learn to respond to a cluster of input population activity would necessarily develop new feature selectivity without requiring any kind of supervision. Because of this property, cluster learning can potentially be repeated within the cortical hierarchy (Fig. 1d).

Here, we simulated a biologically plausible neural circuit that is capable of on-line learning of such clusters from measured neural population data. We found that a winner-take-all feedforward network can, indeed, successfully learn and then readout clusters of real neural activity.

## RESULTS

### A downstream circuit model whose readouts develop strong cluster tuning

In prior work, we described the probability distribution over neural activity patterns with complex models, such as a hidden Markov model, and used machine learning methods to fit the parameters of these models [12]. While this analysis provided insight into the nature of the population code, downstream neural circuits are almost certainly not able to use the same algorithm. In particular, the machine learning techniques involved multiple, *offline* passes through the entire data set. Furthermore, the resulting model had many parameters, and it is unclear how the brain might encode all of these parameters.

We thus sought to investigate an in *silico* simulation of a downstream neural circuit capable of extracting the neural activity clusters via *online*, biologically plausible learning dynamics. This network only seeks to learn clusters in its neural population input, not infer all of the parameters of machine learning models. Importantly, we performed this task on the measured activity of populations of retinal ganglion cells, neurons which serve as the visual input to the cortex.

Based on the theory developed in [16–18], our network consisted of a single layer of readout neurons (indexed *k* = 1, ⋯, *M* in Fig. 1c), which receive spikes from afferent sensory neurons (indexed *i* = 1, ⋯, *N*, which correspond to real neurons in our case) via feedforward connections with synaptic weights, *W_ki_*. Winner-take-all (WTA) competition was induced among readout neurons in the downstream circuit via lateral inhibition (denoted *I* in Fig. 1c). This lateral inhibition was common to all readout neurons, and can thus be viewed as a global inhibitory signal that controls the total gain of the circuit. The WTA circuit’s parameters consisted of the set of feedforward synaptic weights, {*W_ki_*}, and a set of intrinsic excitability parameters, {*b_k_*}, for each readout neuron *k*. The learning dynamics changed synaptic weights in a Hebbian fashion and changed readout neuron biases in a homeostatic fashion (see Eq. 9 of Methods). These changes were made online, i.e. they occurred every 20 ms after the presentation of a single population input pattern (see Methods for numerical simulation and op timization details). Importantly, these plasticity rules were local.

After learning reached a steady-state, individual readout neurons in this WTA circuit developed strong tuning for individual clusters of neural activity in the input (Fig. 2a *bottom*). In contrast, these same readout neurons exhibited no cluster tuning prior to learning (Fig. 2a *top*). Learning occurred quickly, i.e. in less time than required to sample every neural activity pattern (Fig. 2b). Moreover, we found that when the number of readout neurons in the simulated downstream circuit was set equal to the number of known neural activity clusters, the circuit developed a nearly one-to-one correspondence between readout neurons and neural activity clusters at the population level (Fig. 2c; *Dataset* #1). Note that each row of the confusion matrix in Fig. 2c corresponds to one tuning curve for a WTA readout neuron (after learning). Perfect agreement between the activity of readout neurons and the clusters inferred by the machine learning algorithm would result in a confusion matrix with non-zero elements only on the diagonal.

**FIG. 2:**
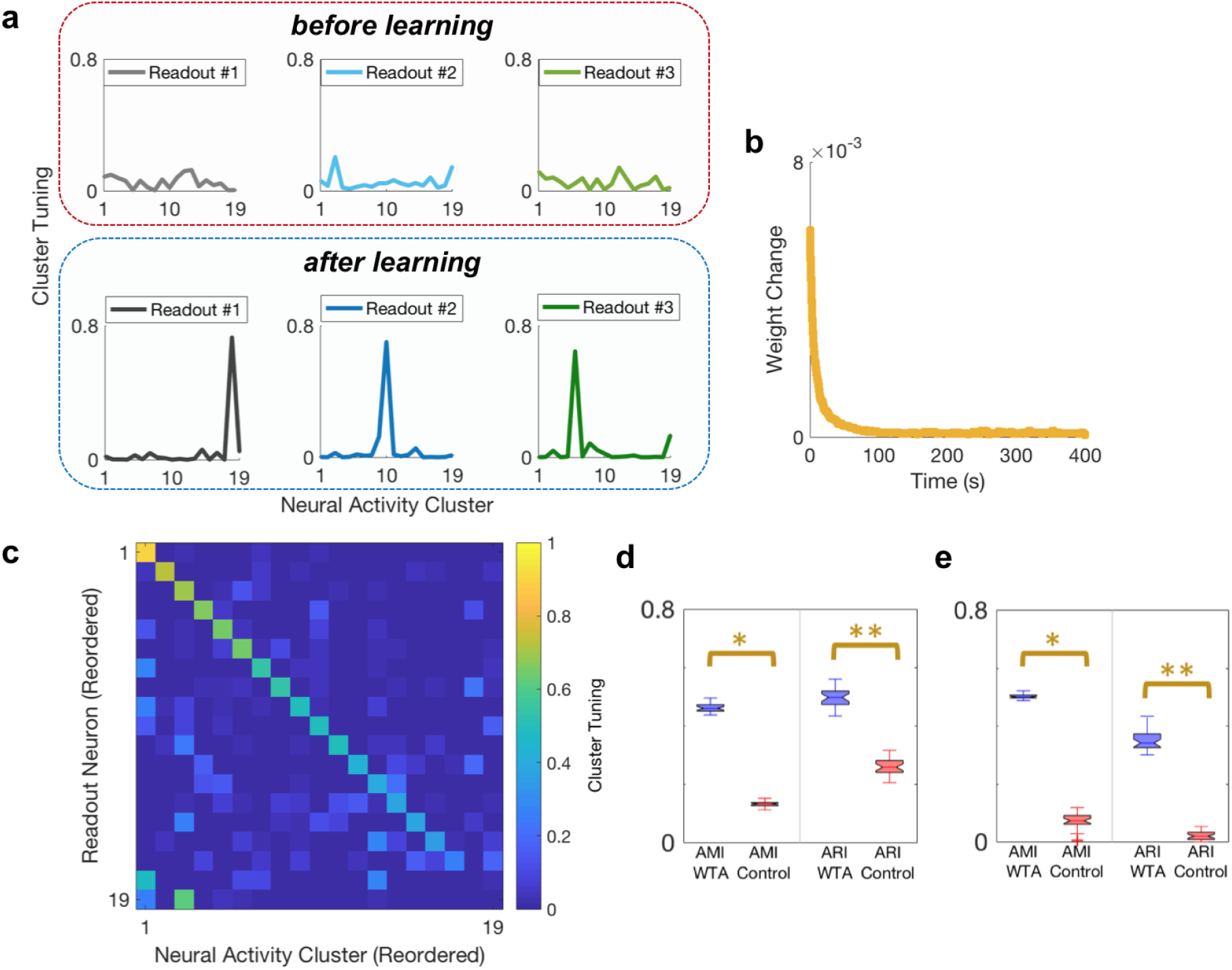
Development of cluster tuning. **(a)** Example tuning curves for three readout neurons of the simulated downstream WTA network before learning (*top*) vs. after learning (*bottom*) on the afferent population activity in *Dataset* #1 (see Methods). Each panel plots the spiking probability of readout neuron, *k*, as a function of the neural activity cluster, *α*, present in the input, *p*(*k*|*α*). **(b)** Mean absolute synaptic weight change, 〈|Δ*W_ki_*(*t*)|〉, as a function of time during learning; curve has been boxcar smoothed (window = 200 time bins). **(c)** Confusion matrix for the WTA circuit after learning on *Dataset* #1, with *M* = 19 readout neurons. Each entry (*k, α*) of the confusion matrix denotes the probability, averaged over all time bins, that readout neuron *k* spiked when the afferent population activity was a member of cluster α. The indices have been reordered so that the confusion matrix is in diagonal form, with the highest values in descending order. Note that perfect decoding performance would correspond to a confusion matrix with a value of one along the diagonal, and zero everywhere else. **(d,e)** Cluster tuning of the WTA circuit after learning (blue) is significantly better compared to a control network (red, see text and Methods) for *Dataset* #1 and #2, respectively.

We quantified cluster tuning by calculating two metrics: the adjusted mutual information (AMI) and the adjusted rand index (ARI), which compare the similarity of two clusterings of the same set of elements [19] (see Methods for details). Both measures range from 1, when there is exact agreement, down to 0 or an expected value of 0, when there is only chance agreement. For *Dataset* #1 and *M* = 19 readout neurons, we found that the AMI and ARI were 0.46 ± 0.015 and 0.50 ± 0.032, respectively (mean ± one standard deviation over 30 realizations of the WTA circuit; Fig. 2d, blue). For *Dataset* #2 and *M* = 50 readout neurons, we obtained similar AMI and ARI values of 0.52 ± 0.010 and 0.35 ± 0.037, respectively (Fig. 2e, blue). In both cases, the clustering was highly significant compared to the level of clustering present before learning (one-sided Mann-Whitney *U* test; *p* =1.5 × 10^-11^) (Fig. 2d,e, red). This control indicates that simply forming many-onto-one mappings from neural activity patterns onto clusters does not, by itself, induce cluster tuning. Instead, Hebbian plasticity appears crucial to shape the distribution of synaptic weights to separate and identify clusters.

### Readout neurons exhibit error correction

Previous work has demonstrated that clusters of retinal ganglion cell activity exhibit error-correction – namely, highly reliable activation by the same visual stimulus [12]. Thus, we wanted to test whether the activity of readout neurons in the WTA circuit could similarity exhibit error correction. Specifically, we implemented WTA learning dynamics on retinal ganglion cell populations responding to a non-repeated natural movie stimulus, and then assessed error correction performance over an interleaved repeated stimulus segment (see Methods). We found that a highly variable set of distinct neural activity patterns were typically elicited at a given time during the natural movie (Fig. 3a). In fact, the detailed activity pattern was often different on each of the 73 repeated trials tested (Fig. 3b *bottom*). However, individual readout neurons could be reliably activated on a large faction of all repeated trials (Fig. 3b *top*), thus exhibiting error correction.

**FIG. 3:**
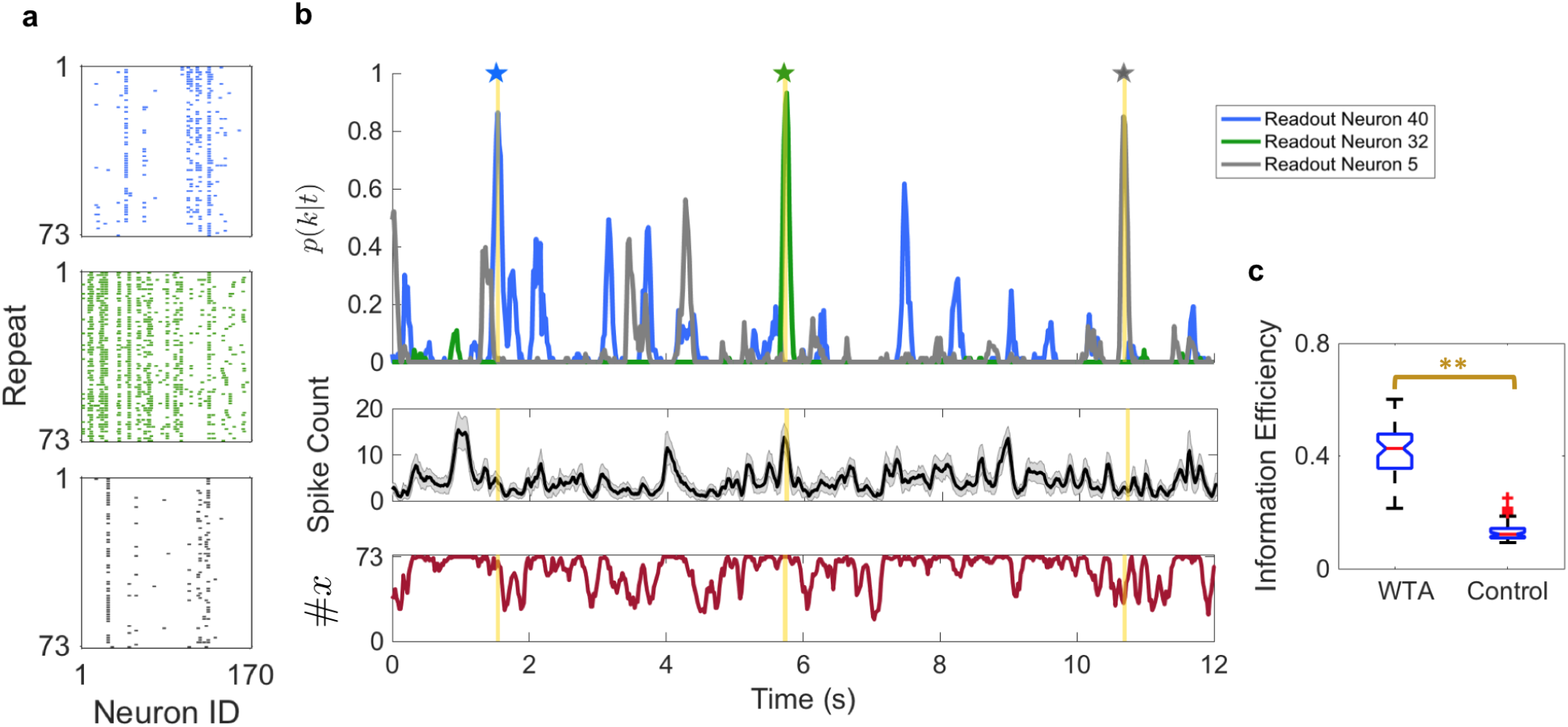
Downstream readout neurons inherit error correcting properties. **(a)** Example spike rasters of the input (retinal) population activity patterns observed over 73 repeats of the 60-sec target movie segment for *Dataset* #2. Examples are shown for each of three time bins in the target movie segment, which are highlighted in yellow in panel (b). Each row is the observed input population activity pattern on one trial of the stimulus. White denotes silence; colors denote spikes from each of three readout neurons. **(b)** *Top:* Spiking probability of each of three readout neurons (denoted by color) plotted versus time within the natural movie. *Middle*: Mean spike count of the input population activity pattern, averaged over the repeats. Error bars denote one standard deviation. *Bottom*: The number of distinct input population activity patterns observed over all 73 repeats at each time. **(c)** Information efficiency of the WTA circuit vs. the random partition control (see Methods).

To quantify the reliability of the readout neurons, we calculated the information efficiency (see Methods). Information efficiency values for simulations of the WTA circuit (median 0.43) were significantly greater than those of matched random partition controls (Fig. 3d; *p* < 1 × 10^-100^, Mann-Whitney *U* test; see Methods). This control analysis indicates that the error correction exhibited by readout neurons in the WTA circuit was not trivially due to compressing tens-of-thousands of input activity patterns into a much small number of clusters. Rather, learning the correct neural activity clusters was crucial.

### Learning all of the input clusters

While the WTA circuit developed strong tuning for most clusters, some other clusters were not efficiently captured (see Fig. 2c, bottom right). We hypothesized that adding more readout neurons to the WTA circuit would allow for the detection of all input clusters. To this end, we used *Dataset* #2 as the input, which consisted of *m* = 50 measured neural activity clusters, and formulated a WTA circuit with twice as many readout neurons (*M* = 100). We found that the 2-fold redundant WTA circuit developed complete tuning for the entire set of clusters (Fig. 4b), whereas a WTA circuit without redundancy (*M* = 50) developed tuning to only ~ 75% of the clusters (Fig. 4a). Moreover, many of the readout neurons in the redundant WTA circuit developed strong tuning for two or three neural activity clusters (Fig. 4c, d). This mixed selectivity for input neural clusters was not present when the number of readout neurons matched the number of clusters. Thus, a larger layer of readout neurons can potentially confer the additional advantage of mixed selectivity tuning.

**FIG. 4:**
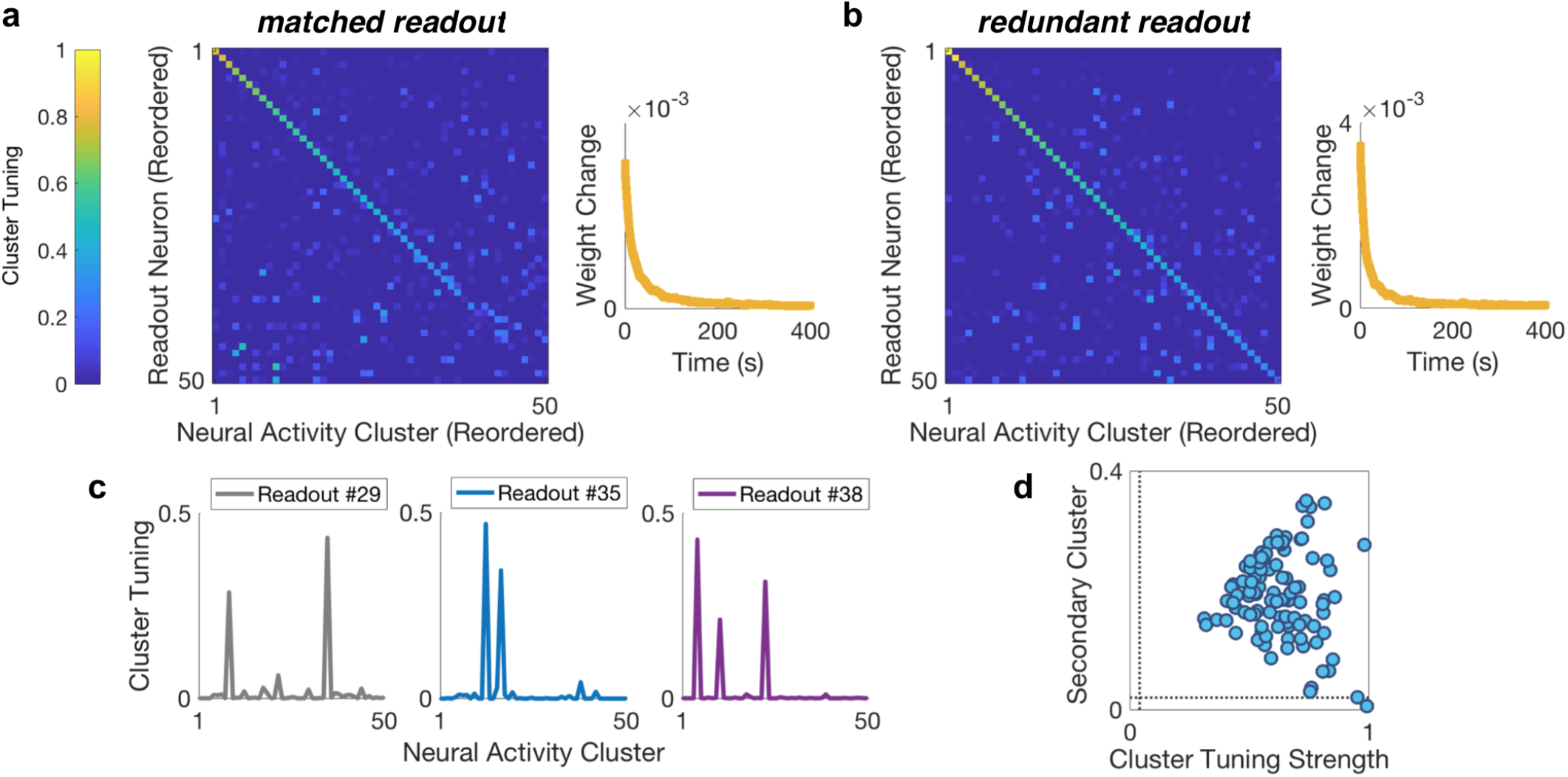
Redundant WTA circuits and mixed cluster selectivity. **(a,b)** *Left*: Confusion matrix for the WTA circuit after learning on the ganglion cell population activity in *Dataset* #2, when the number of WTA readout neurons was set to either (a) *M* = 50 or (b) *M* = 100. *Right*: Mean absolute synaptic weight change as a function of time during learning. **(c)** Cluster tuning curves for three example readout neurons in the redundant WTA circuit. **(d)** Scatter plot of the magnitude of the second largest peak in the tuning curve (y-axis) vs. the combined magnitude of the maximum and second largest peak (x-axis); see Methods for definitions. Each blue dot represents a single readout neuron. Note that uniformly random tuning curves would be located at the intersection of the dashed grey lines.

## DISCUSSION

In summary, we have constructed a simple, biologically plausible decoding circuit that takes as an input the spiking activity of a population of neurons and learns the clusters of activity present in that population. The outcome of the learning process is that individual readout neurons in the decoding circuit develop strong cluster tuning, and different readout neurons specialize for different clusters in the input population. The basic ingredients of this circuit are: 1) Hebbian plasticity in the feedforward synapses, 2) winner-take-all competition among readout neurons, and 3) homeostatic plasticity of the excitability of readout neurons. Hebbian plasticity is, in some sense, the core operation that differentiates the pattern of neural activity in different input clusters. Winner-take-all inhibition causes different readout neurons to specialize for different input clusters. Finally, homeostatic plasticity of excitability helps to encode the probability of each cluster occurring.

While the WTA circuit contains computational elements and learning rules that are broadly present within the central brain, it is obviously not a detailed model of any one brain area. We expect that additional realism would impact the learning rate and might also diminish the overall performance. However, it is important to keep in mind that the goal of the readout circuit might not be to achieve perfect cluster tuning; in fact, we suspect that developing mixed tuning for several related clusters might be a more robust solution. In any case, our results clearly show that the WTA circuit develops significant cluster tuning through its learning dynamics.

A key feature of our approach has been to use measured rather than simulated spike trains as the input to the circuit. This test is more stringent and relevant than previous analyses. Mathematically, the WTA circuit stochastically learns a mixture model [17, 18], which is a simpler, special case of the Tree hidden Markov model that best fits retinal ganglion cell population activity (see Methods). Because of this difference, the ability of the WTA circuit to learn clusters from these real spike trains was not guaranteed. Furthermore, the result suggests that this type of WTA mechanism might also be able to learn clusters from population neural activity elsewhere in the brain that can be described by a wider class of Markov models.

Because of the simplicity of the WTA circuit, it is plausible that such an operation might be carried out in multiple brain areas. One proposal is that the WTA circuit is implemented in layer 4 of the cortical microcircuit (Fig. 1d). Then, when signals flow into layer 2/3, they are recombined across larger spatial scales, creating new correlations among neurons that form a population code with a new set of clusters. As a result, when these signals flow up the cortical hierarchy, the next layer 4 can again implement a WTA circuit to learn these new clusters. Because the receptive fields represented by clusters and individual neurons differ [12], layer 4 readout neurons will represent receptive field properties that are qualitatively new compared to the input neurons. This property agrees with the general structure of the visual hierarchy, in which a new feature basis set emerges in layer 4 and is maintained throughout the non-granular layers.

In addition, this hypothesis makes a prediction about pairwise correlations in the cortex. Fundamentally, it is the correlation among neurons on a fast time scale that defines clusters of population activity. When the WTA circuit learns these clusters, its readout neurons will then have relatively low correlation. And when signals are recombined over large spatial receptive fields in layer 2/3, correlations will increase, reflecting correlations in the visual scene present over larger spatial scales. Thus, this hypothesis predicts smaller correlation in layer 4 than non-granular layers, as seen experimentally [20]. In this context, it will be interesting to test whether clusters are present in layer 2/3 of V1 and whether the receptive fields of these clusters match those of layer 4 neurons in extrastriate cortex.

## MATERIALS AND METHODS

### Ethics statement

This study was performed in strict accordance with the recommendations in the Guide for the Care and Use of Laboratory Animals of the National Institutes of Health. The protocol was approved by the Institutional Animal Care and Use Committee (IACUC) of Princeton University (Protocol 1828).

### Experimental procedures

#### Electrophysiology

The input activity patterns provided to the simulated downstream circuit model were obtained from electrophysiological recordings of larval tiger salamander (*Ambystoma tigrinum*) retinal ganglion cells responding to naturalistic movies. Detailed experimental procedures are reported in [11, 12]. In brief, animals were euthanized according to institutional animal care standards. Experiments were conducted with the pigment epithelium intact and surgery performed in dim light. After transferring to oxygenated Ringer’s medium at room temperature, the isolated retina was lowered with the ganglion cell side against a custom 252-electrode multi-electrode array (MEA). Raw voltage traces were digitized and stored for offline analysis using a 252-channel preamplifier (MultiChannel Systems, Germany). Offline spike sorting was performed using custom software. Detailed recording and analysis methods, including the spike-sorting algorithm, are described in [21]. For the two natural movie experiments reported here, 152 and 170 neurons passed the standard tests for waveform stability and lack of refractory period violations, respectively.

#### Visual stimulus display

Stimuli were projected onto the multi-electrode array from a CRT monitor at a frame rate of 60 Hz, and gamma corrected for the display. The natural movie stimulus consisted of a 7-min gray scale recording of leaves and branches blowing in the wind. We conducted the natural movie experiment with two different designs, yielding two datasets: (1) in the first, which we refer to as *Dataset* #1, the movie was looped five times, but with a different pixel centered on the recording area for each repeat; (2) in the second, which we refer to as *Dataset* #2, we interleaved 73 unique movie segments, each 60 s in duration, with 73 repeats of a fixed 60-s fixed movie segment. Design (1) was constructed this way to provide a non-repeated stimulus: since the patch of retina recorded by the multi-electrode array subtended only a small portion of the stimulus, the retinal input was effectively non-repeated over the full recording session. *Dataset* #1 was approximately 30 min in duration, while *Dataset* #2 was approximately 2 hours and 26 min in duration.

### Data preparation

For each dataset, we discretized spike trains into 20 ms time bins, as in previous work [11–13, 22, 23]. This yielded a sequence of binary population response vectors, which we denote as 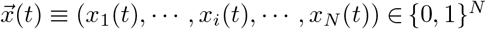, where *i* = 1, ⋯, *N* labels the neuron identity and *t* the time bin. We set *x_i_*(*t*) = 1 if neuron i fired at least one spike within time bin *t*, and 0 otherwise.

### Mathematical details for the network model

#### Downstream network architecture

For the readout neurons of the downstream circuit model (which was numerically simulated, as detailed in the following section), a stochastic model was used similar to that proposed in [16–18]. Specifically, the instantaneous spiking probability of readout neuron k during time bin *t* (for Δ*t* → 0), here denoted *ρ_k_*(*t*), is characterized by an exponential dependence on its current membrane potential, *v_k_* (*t*), and a common, time-varying inhibitory contribution *I*(*t*):

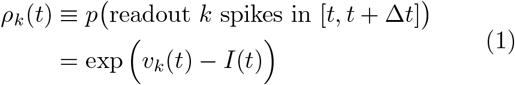

The membrane potential of readout neuron *k* of the WTA network is given by

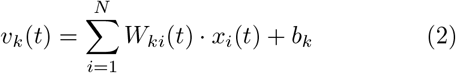

where *b_k_* formalizes neuron *k*’s local intrinsic excitability bias (see next sections for details).

For a given time bin *t*, each readout neuron receives the same global inhibitory contribution *I*(*t*), given by

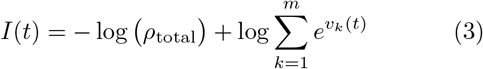

where *ρ*_total_ denotes the total number of spikes over all WTA readout neurons per time bin. Note that *ρ*_total_ is a hyperparameter that is constant in time. In practice, we used Δ*t* = 20 ms per time bin, and *ρ*_total_ = 1 spike. This corresponds to only allowing one WTA readout neuron to spike per time bin, i.e. strict winner-take-all. It is this lateral inhibition term in Eq. 3 which serves to introduce competition among the readout neurons.

#### Implicit statistical model of the upstream neural population code

Our choice of the circuit architecture and update rules for the network parameters was motivated by the connection to a simpler, special case of the full machine learning model that we used in our previous work in [12] to identify the noise-robust clusters from retinal ganglion cell population data. This simplified model assumes that time bins are statistically independent, and also lacks the tree-based interneuronal correlations included in the full model. Although the full model performs better, this simplified model still fits the retinal ganglion cell population data reasonably well (see the Supplement in [12]). Formally, this scenario corresponds to modeling the input retinal ganglion cell population activity patterns as being generated by a Bernoulli mixture model, which models the joint probability of the afferent population activity as:

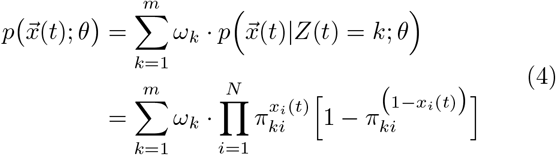

where *Z*(*t*) is a random variable that represents the latent cluster in which the input population activity pattern 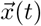 resides in population activity space (or equivalently, codeword), *m* denotes the total number of neural activity clusters, and *ω_k_* denotes the mixing weight of cluster *k*.

Applying the theoretical approach developed by [16–18], the WTA downstream network model parameters can be explicitly related to the Bernoulli mixture model parameters *θ* ≡ ({*π_ki_*}, {*ω*_*k*_}) by:

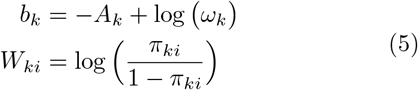

where

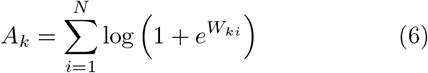

Using this correspondence in Eq. 5, and provided that certain assumptions are met (see next sections), [18] showed that the WTA circuit model can represent an internal, implicit Bernoulli mixture model.

### Numerical simulation of the downstream circuit

All computer simulations of the downstream circuit described above were implemented using custom code written in C++, combined with Mex files for compatibility with post-processing in Matlab. The code used to generate the results presented here will be freely available as a repository on Github, which can be accessed at: https://github.com/adriannaloback.

We used as input to our numerically simulated downstream circuit the sequence of spike times observed in the actual retinal population data. The input is defined as *x_i_*(*t*) = 1 if input neuron *i* emitted a spike within the last Δ*t* ms, and 0 otherwise, corresponding to a rectangular post-synaptic potential (PSP) of length Δ*t*. I.e. at each discrete time step *t*, the input to our simulated downstream network was a population activity pattern 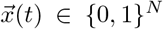. It is important to emphasize that all afferent neural activity patterns on which learning dynamics were performed were obtained from actual retinal population data. For all results presented here, a time bin width of Δ*t* = 20 ms and *ρ*_total_ = 1 spike was used.

For each simulation, we used constant learning rate hyperparameters *η_b_*, *η_W_*, which were chosen via a grid search. When *Dataset* #1 was provided as the afferent input population activity, we found via this grid search approach that the optimal learning rate values were (*η_b_* = 0.1, *η_W_* = 0.25) for *M* = 19 readout neurons, and (*η_b_* = 0.05, *η_W_* = 0.25) for *M* = 38 readout neurons. When *Dataset* #2 was provided as the input, the optimal learning rate hyperparameter values identified via the grid search were (*η_b_* = 0.25, *η_W_* = 0.25) and (*η_b_* = 0.75, *η_W_* = 0.75) for *M* = 50 and *M* = 100 readout neurons, respectively.

For the results presented in Figs. 2 and 3, the number of readout neurons *M* in the simulated downstream WTA network was set equal to the number of latent modes, m, that were identified via 2-fold cross-validation when fitting the Tree hidden Markov model to the same respective dataset of afferent retinal population responses (*m* = 19 for *Dataset* #1 and *m* = 50 for *Dataset* #2). For the redundant WTA circuit in Fig. 4, twice as many readout neurons were used than the number of neural activity clusters identified by the Tree hidden Markov model.

#### Initialization of network parameters

It is important to choose distinct initial *W_ki_* terms for each downstream WTA readout neuron *k*, because if two readout neurons have identical initial parameters, the learning dynamics will not separate them. We therefore generated the *π*_*k*_(0) ≡ (*π*_*k*1_(0), ⋯, *π_kN_*(0)) vectors randomly and independently from the uniform distribution over [0.45,0.55]. The initial synaptic weights were then set to *W_ki_*(0) = log (*π_ki_*(0)/(1 − *π_ki_*(0))). The target *μ_k_* values (see next section) used when implementing Eq. 9 were chosen to match the mixing weights of the fit Tree hidden Markov model for the respective input dataset, i.e. we set *μ_k_* = *ω_k_*. Each *b_k_* network parameter was then initialized to *b_k_*(0) = – *A_k_*(0) + log(*μ_k_*).

### Hebbian and homeostatic plasticity rules

#### The need for homeostatic plasticity

In addition to the spiking input, each readout neuron’s membrane potential *v_k_* features (see Eq. 2) an intrinsic excitability *b_k_*. Besides the prior constant log(*ω_k_*), this intrinsic excitability depends on the normalizing term *A_k_* (see Eq. 5), and consequently on all afferent synaptic weights through Eq. 6. In the presence of synaptic plasticity, i.e. time-varying weights *W_ki_*, it is unclear how biologically realistic neurons could communicate ongoing changes in synaptic weights from distal synaptic sites to the soma. Thus, for the purposes of a biologically plausible downstream circuit, we cannot assume that readout neurons could compute *A_k_* directly.

This poses a challenge in terms of the mathematical theory, since the theory requires that the WTA readout neurons must be able to sample from the correct posterior distribution 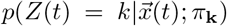 of the implicit model given by Eq. 4 (where *π*_k_ denotes the vector 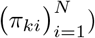 [18]. That is, formally, the theory requires that

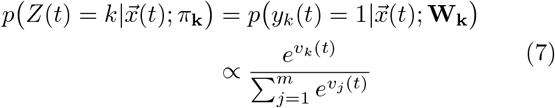

where **W**_k_ denotes the vector 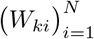; *y_k_*(*t*) denotes the spiking response of readout neuron *k* (i.e. *yk*(*t*) = 1 if readout neuron *k* spiked during time bin *t*, and is 0 otherwise); and the second line of Eq. 7 follows from Eq. 1 and Eq. 3. Letting 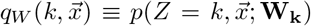 and applying Bayes’ theorem, we see that this posterior of the implicit model is given by:

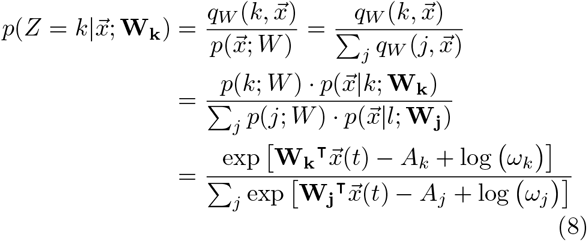

Thus, in general the posterior problematically involves computing the *A_k_* terms, but requiring readout neurons to compute *A_k_* is not biologically plausible. To circumvent this critical issue, [18] assumed uniform values of *A_k_* across all readout neurons, as well as uniform mixing weight *ω_k_* values. Note that these two assumptions allow for canceling out (and hence no longer needing to compute) the *A_k_* terms in Eq. 8, while also satisfying Eq. 7.

However, the assumption of uniform *A_k_* and mixing weight *ω_k_* values was grossly violated for our afferent population activity data, so we could not use the trick used in [18]. Thus, to cope with this issue while maintaining biological plausibility during learning, we used the homeostatic plasticity approach introduced in [17]. This approach applies homeostatic dynamics to the intrinsic excitabilities in a manner that eliminates the non-local terms *A_k_* terms from the network dynamics. Using this approach, once the intrinsic excitabilities *b_k_* were initialized (see previous section), they were then fully regulated by the local homeostatic plasticity rule given in the first line of Eq. 9, which requires only local knowledge.

#### Rules for the learning dynamics

In practice, we implemented the following plasticity rules for the learning dynamics in the downstream circuit:

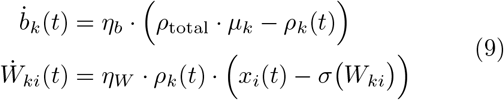

where *η_b_* and *η_W_* are learning rate hyperparameters, *x_i_*(*t*) is the response of afferent (input) neuron *i*, 0 < *μ_k_* < 1 denotes a target average activation for WTA readout neuron *k* that the homeostatic plasticity will aim to maintain, and *σ*(*x*) = (1 + exp(−*x*))^−1^ denotes the logistic function. Importantly, these updates were performed on a per-sample basis (i.e. for each input population activity pattern), and could thus be implemented in real time.

As derived in [17], these plasticity rules instantiate a sample-based, stochastic version of the variational Expectation Maximization scheme for learning the network parameters that maximize the likelihood of the internal latent cause model of the upstream population activity (Eq. 4). Note that we are not interested in the network parameters per se, but rather the corresponding learned clusters underlying the internal model.

### Fitting the Tree hidden Markov model

The Tree hidden Markov model developed in [12] models the sequence of observed afferent population responses as:

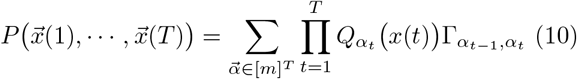

where *T* is the total number of time bins, 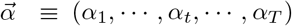 is the sequence of hidden cluster identities, and *Q_α_*(·) denotes the emission distribution for cluster *α* ∈ [*m*] ≡ {1, ⋯, *m*}. The transition matrix Γ, which has entries Γ_*α*_*t*−1_,*α_t_*_ ≡ *P*(*α_t_*|*α*_*t*-1_), is assumed to be stationary (i.e. time-independent). Each emission distribution *Q_α_* in the model is a tree graphical model (see [24]).

For a fixed number of latent modes m, the model was fit to data using the same learning procedure presented in [12]. In brief, the model parameters were inferred by maximum likelihood, using the Baum-Welch algorithm with an M-step modified to accommodate the tree graphical model form of the emission distributions *Q_α_*. Full details of the algorithm are described in [12]. The code used to fit the model was written in C++.

To select the number of latent modes m, also called the “latent dimensionality” [25], we carried out a 2-fold cross-validation procedure in which we randomly chose half of the time bins in the data to assign to the training set (with the other non-overlapping half assigned to the test set). The latent dimensionality was then chosen to be the value that maximized the cross-validated log likelihood (CV-LL), averaged over the 2 folds. Note that to mitigate overfitting, we also incorporated a regularization parameter *λ* ∈ [0,1] in this procedure (in practice, *λ* = 0.002 was used throughout), as in [12].

To obtain the mixing weight terms *ω_k_* fit by the Tree hidden Markov model, note that we used the stationary distribution 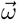 of the fit Markov chain. Formally, 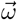 is defined as the left eigenvector of the transition matrix Γ with unity eigenvalue (i.e. 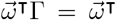) and satisfies 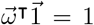. The corresponding stationary distribution of the afferent population activity, for a single population response 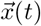, is then modeled as:

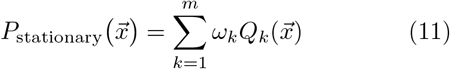

In order to compare the clustering scheme performed by the simulated WTA circuit to that of the (non-biologically plausible) Tree hidden Markov model (see next section), we first computed the clustering induced by the fit Tree hidden Markov model on each sequence of afferent activity data. To compute this clustering, we used the Viterbi algorithm, which is a standard method that computes the most likely (maximum *a posteriori*) sequence of neural activity clusters (i.e. latent modes of the hidden Markov model).

### Quantifying cluster tuning

To compare how the simulated downstream circuit (after learning) clustered afferent neural population activity patterns relative to the machine learning approach used in [12], we applied two standard metrics: (1) the adjusted mutual information (AMI), and (2) the adjusted rand index (ARI). Both quantify the similarity between clustering schemes implemented on the same dataset 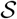. In our case, 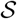 is the set of observed input population activity patterns, i.e. 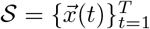.

Let *C* = {*C*_1_, ⋯, *C_M_*} denote the set of clusters induced by the downstream WTA network, i.e. each *C_k_* is a cluster of afferent population activity patterns that readout neuron *k* learns to develop tuning specificity for (i.e. readout neuron *k* will spike selectively when an activity pattern from cluster *C_k_* is presented as input). Similarly, let 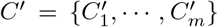 denote the set of neural activity clusters identified by the Tree hidden Markov model. The *normalized confusion matrix* Ψ of the pair *C,C′* is the *M* × *m* matrix whose *k*, *α*-th entry equals the number of elements in the intersection of the clusters *C_k_* and 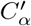:

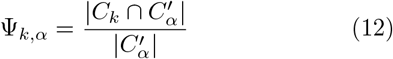

where 1 ≤ *k* ≤ *M*, 1 ≤ *α* ≤ *m*. Note that the confusion matrices shown in Fig. 2 and Fig. 4 are exactly the normalized confusion matrices of the two clustering schemes, Ψ, defined in Eq. 12.

The AMI is defined as:

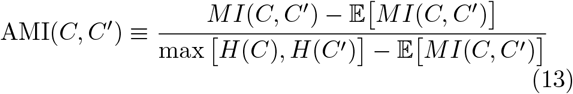

where

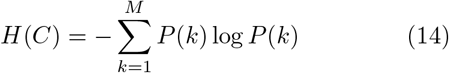

is the entropy of the clustering scheme, where

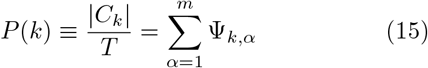

and *MI*(*C, C′*) denotes the mutual information between the two clustering schemes. The AMI has value 1 when the normalized confusion matrix Ψ has weight only on the diagonal, and is 0 when the mutual information equals the value expected due to chance alone.

Similarly, the adjusted rand index (ARI) is the corrected-for-chance version of the Rand index [19], formally given by:

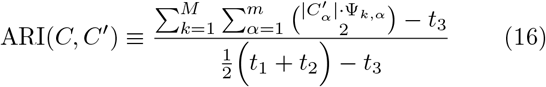

where

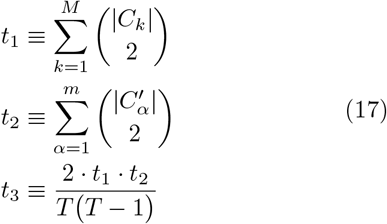

The ARI has a value of 1 when the two partitions are identical, and its expected value is 0 in the case of random clusters.

### Random network controls

For the random network control analyses (see main text and Fig. 2d,e), we simulated an analogous version of the downstream WTA network, in which no learning was implemented. That is, the control network was initialized with random *W_ki_* parameter values as in the actual network, but no learning dynamics were implemented to update these random initialized weights. It is important to note however that the control network was provided with the ‘correct’ target threshold values *μ_k_* (which were set equal to the mixing weight parameter values, *ω_k_*, identified by fitting the Tree hidden Markov model), as this was also provided to the actual downstream WTA network (i.e. with learning). The same number of readout neurons was used in each control network as the respective true WTA network.

### Statistical Tests

Reported *p* values for the comparison of the AMI and ARI distributions for the simulated control network versus the true downstream WTA network (i.e. with learning) were calculated using the one-sided Mann-Whitney *U* Test.

### Information efficiency

As in [12], to compute the information efficiency of the simulated WTA circuit after learning (Fig. 3c), for each readout neuron we computed the fraction of stimulus repeats (for Dataset #2) on which that readout neuron was active at time bin *t*, denoted *r*(*t*), and its time average, 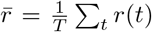. We then defined the output entropy for the readout neuron, *S*_out_, and its noise entropy, *S*_noise_, as follows:

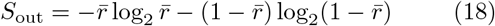

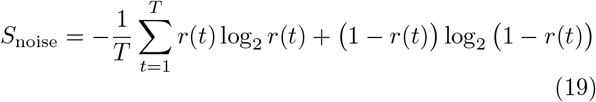

The information efficiency was then defined to be the mutual information divided by the output entropy:

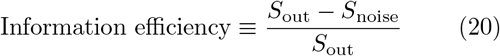

### Random partition control

This control was based on constructing a deterministic mapping from afferent population activity patterns to downstream WTA readout neurons. The goal was to partition input patterns into a set of randomly assigned readout neurons with number and probability distribution matched to that of the actual downstream WTA circuit after learning. To do this, we first assigned each readout neuron of the control circuit a “capacity” initially equal to the probability of the true WTA circuit. We then ranked the unique input patterns in decreasing order of probability. Proceeding in rank order, we mapped each unique input pattern to a random control readout chosen from the set of readouts with capacity greater than the pattern’s probability. After assigning an input pattern to a control readout, that control readout neuron’s capacity was decremented by the input pattern’s probability before assigning the next input pattern. Eventually, input patterns were encountered with probability greater than any control readout neuron’s remaining capacity. In this case, the input pattern was simply assigned to a random downstream control readout unit. Since the probability values of these input patterns were tiny, the probability distribution of the randomly constructed control readout units was extremely close to that of the true readout units of the true downstream WTA circuit model after learning.

### Quantifying mixed selectivity

To visualize the distribution of single vs. mixed cluster selectivity over the population of readout neurons of the redundant WTA circuit (*M* = 100 readout neurons; see Fig. 4d), we computed two quantities for each readout neuron, as described below. For a given readout neuron *k*, let 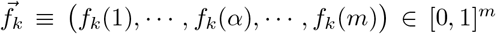 denote its tuning curve (after learning) over all *m* neural activity clusters. That is, *f_k_*(*α*) represents the empirical probability (after learning reached steady state) of readout neuron *k* firing a spike when neural activity cluster *α* was active.

For each tuning curve, we computed the magnitude of the second largest peak, *ν*_2_ ≡ max_*α*∈[*m*]\*α^*^*_ *f_k_*(*α*), where *α*^*^ denotes the neural activity cluster corresponding to the maximal peak. We also computed the combined magnitude of the maximal and second largest peak, given by *ν*_1_ + *ν*_2_, where *ν*_1_ denotes the magnitude of the maximal peak. Each point in Fig. 4d corresponds to (*ν*_2_, *ν*_1_ + *ν*_2_) for one tuning curve.

## SUPPLEMENTARY TEXT

### Test Case

As a sanity check to ensure that our custom-written code for the downstream WTA circuit simulations was working properly, we simulated a test case “dataset” for which the correct network parameters and clusters was analytically known. To do this, we first used the standard batch Expectation Maximization (EM) algorithm [26] to fit a Bernoulli mixture model with *m* = 13 latent clusters to *Dataset* #1, which models the joint probability of the population activity as:

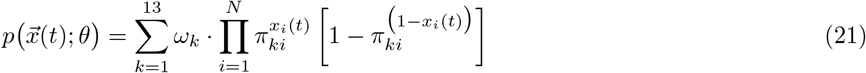

where *N* = 152 afferent retinal ganglion cells. Note that the batch EM algorithm is not biologically plausible - for example, it involves multiple, *offline* passes through the entire data set. (In practice, we implemented 100 offline passes/iterations).

After inferring the maximum likelihood estimated (MLE) parameters via the batch EM algorithm, we then randomly generated *T* = 100, 000 samples 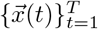 from the fit Bernoulli mixture model (Eq. 21), where each 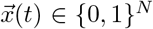 denotes a simulated *N*-dimensional input population activity pattern. Note that for simulating the sequence of input population activity, we used a time bin width of Δ*t* = 20 ms, to match the actual analyses. We then randomly initialized the network parameters of a simulated downstream WTA circuit with *M* = 13 readout neurons. As in the actual analyses, we initialized the intrinsic excitabilities as 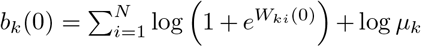, where we set each 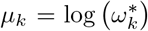, where 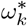 denotes the mixing weight parameter value fit by the batch EM algorithm. Each synaptic weight was randomly initialized as *W_ki_* = log (*π_ki_*(0)/(1 − *π_ki_*(0))), where *π_ki_*(0) was randomly drawn from a uniform distribution over [0.45,0.55].

Since all of the initialized synaptic weights were therefore comparable, prior to learning, the readout neuron with the largest 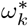 value and hence greatest intrinsic excitability (readout neuron #8) dominated the spiking activity of the WTA circuit (Fig. S1a), as expected. The circuit prior to learning - i.e. without the implementation of the synaptic plasticity dynamics - thus exhibited no cluster tuning. In stark contrast, after learning (i.e., after allowing the network parameters to reach a steady state), all 13 readout neurons exhibited strong tuning selectivity for the known neural activity clusters (Fig. S1b). Note that two of the known clusters corresponded to the all-silent input state; as this state is qualitatively different [11], only the non-silent clusters are shown in Fig. S1c,d. The WTA circuit developed nearly perfect cluster tuning for the 11 known non-silent clusters (Fig. S1c,d).

**FIG. S1:**
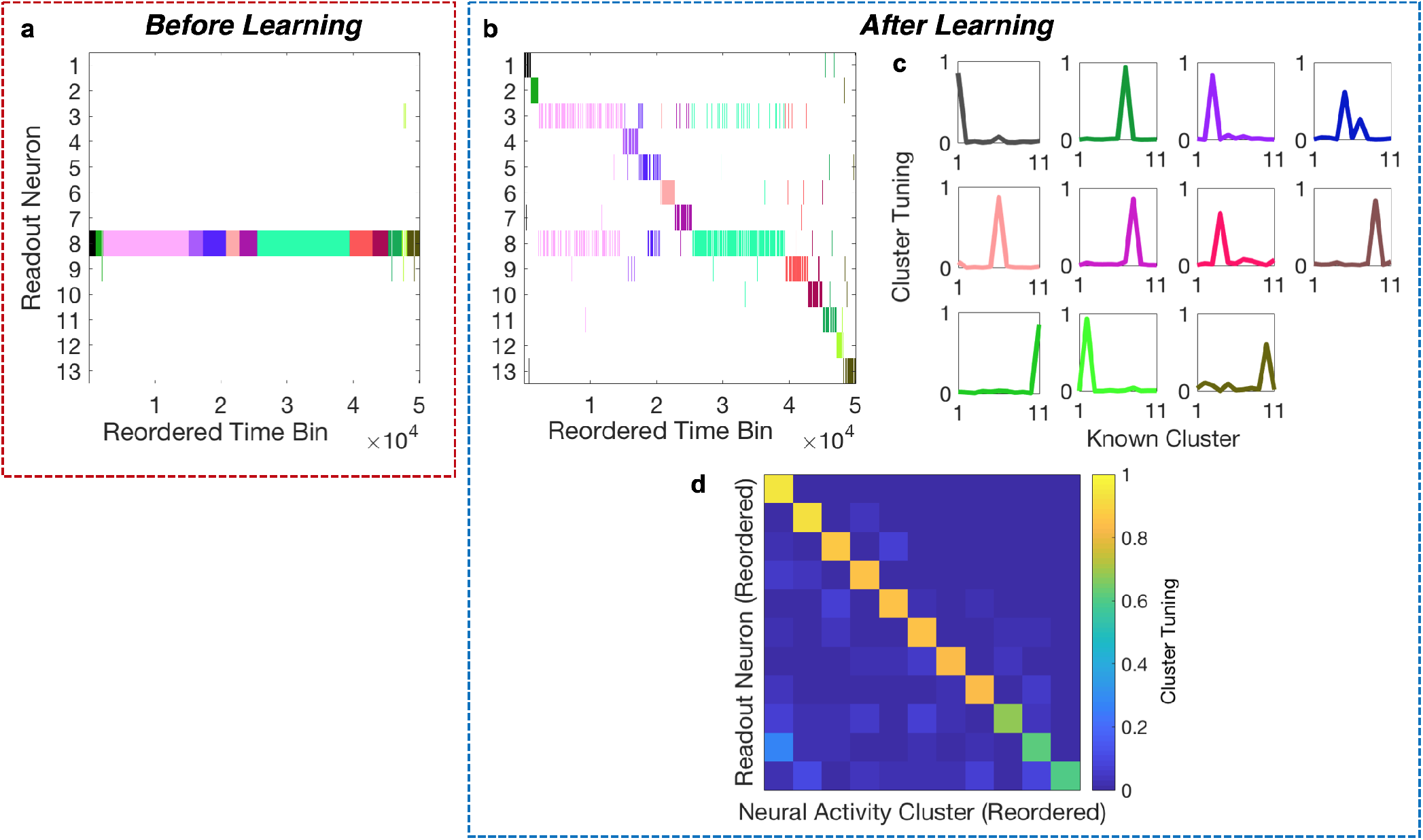
Performance of the WTA circuit on the synthetic test case afferent activity. **(a,b)** Spike raster plots of the WTA circuit (a) prior to learning, i.e. without the implementation of the synaptic plasticity dynamics, vs. (b) after learning. Each row depicts the spike train of one of the 13 readout neurons in the WTA circuit as function of time. Time on the x-axis has been reordered so that time bins during which the 1st known cluster was active are displayed first, time bins during which the 2nd cluster was active are displayed second, etc. Different known neural activity clusters are denoted by different colors. **(c,d)** Shown are (c) the individual tuning curves and (d) corresponding confusion matrix obtained after learning for the 11 readout neurons that developed tuning for the known 11 non-silent clusters (see text). Colors in (c) denote different known clusters and have the same correspondence as in panels (a,b).

